# Adapting ProteinMPNN for antibody design without retraining

**DOI:** 10.1101/2025.05.09.653228

**Authors:** Diego del Alamo, Rahel Frick, Daphne Truan, Joel Karpiak

## Abstract

The neural network ProteinMPNN designs protein sequences capable of folding into predefined tertiary structures and quaternary assemblies. It has become widely used due to its high success rates when working with synthetic topologies rich in secondary structure. Here, we show degraded performance on the complementarity-determining regions (CDRs) of antibodies, with designs frequently failing to resemble native antibodies or failing to refold into the designed conformations. We also show that this underperformance can be rescued by ensembling its predictions with those from the antibody-specific protein language model AbLang, which designs exclusively using sequence information learned from large databases of antibody sequences. Finally, we tested 96 trastuzumab variants with CDRH3 loops redesigned by the ensembled ProteinMPNN+AbLang method and found that it generated thirty-six HER2 binders, compared to three out of 96 designs generated by ProteinMPNN alone. The data highlight the value of incorporating additional restraints derived from language models during structure-based sequence design of antibodies.

## Main text

Over the past three years, the neural network ProteinMPNN^1^ has become the scientific community’s go-to machine learning model for finding amino acid sequences capable of folding into 3D structures of interest^2–6^. Although predominantly applied to topologies rich in secondary structure^3^, anecdotal evidence suggests lower performance on loops, especially longer loops^7^. ProteinMPNN has nevertheless been used to design the complementarity-determining regions (CDRs) of antibodies that mediate antigen binding, with experimentally validated success^8^. In one study, nanobodies with ProteinMPNN-designed CDRs bound their targets with nanomolar affinity and adopted structures matching the design models with sub-angstrom accuracy. However, these hits were derived from large screens with relatively low success rates, and the effectiveness of this method in cases where fewer designs are being evaluated remains unclear.

In this communication, we present computational and experimental evidence that ProteinMPNN struggles to reliably design CDRs with antibody-like sequences that refold into the design pose or bind the target antigens. We also show that this shortcoming can be overcome by ensembling its design predictions with those from AbLang, a protein language model trained exclusively on the sequences of antibody variable regions^9^. The scheme, outlined in Figure S1, does not require retraining of either neural network. AbLang, like other language models trained on a single protein family, exhibits a strong bias toward native-like sequences^10–12^, thus serving as a natural complement for ProteinMPNN, which exclusively relies on structure for inference.

The work presented here was prompted by several observations made when using ProteinMPNN to redesign the CDRs of antigen-bound VH/VL antibodies from a previous benchmark^13^. First and most importantly, the generated sequences were not particularly antibody-like. This was most evident when comparing the designed sequences to those of naturally occurring antibodies that share a similar V-gene, from which most of the variable domain is derived, in the Observed Antibody Space^14^. Comparing CDR1 and CDR2 designs to length- and V-gene-family-matched position-specific scoring matrices (PSSMs) revealed persistent, highly unusual sequence features in ProteinMPNN designs (Figure S2). As a result, the overall designs shared a median of 75% sequence identity with the most similar germline V-genes, far short of the 90% median observed for therapeutic antibodies^15^ (Figure 1A). This shows that structure alone does not sufficiently constrain the antibody design problem toward native-like sequences.

**Figure 1.**
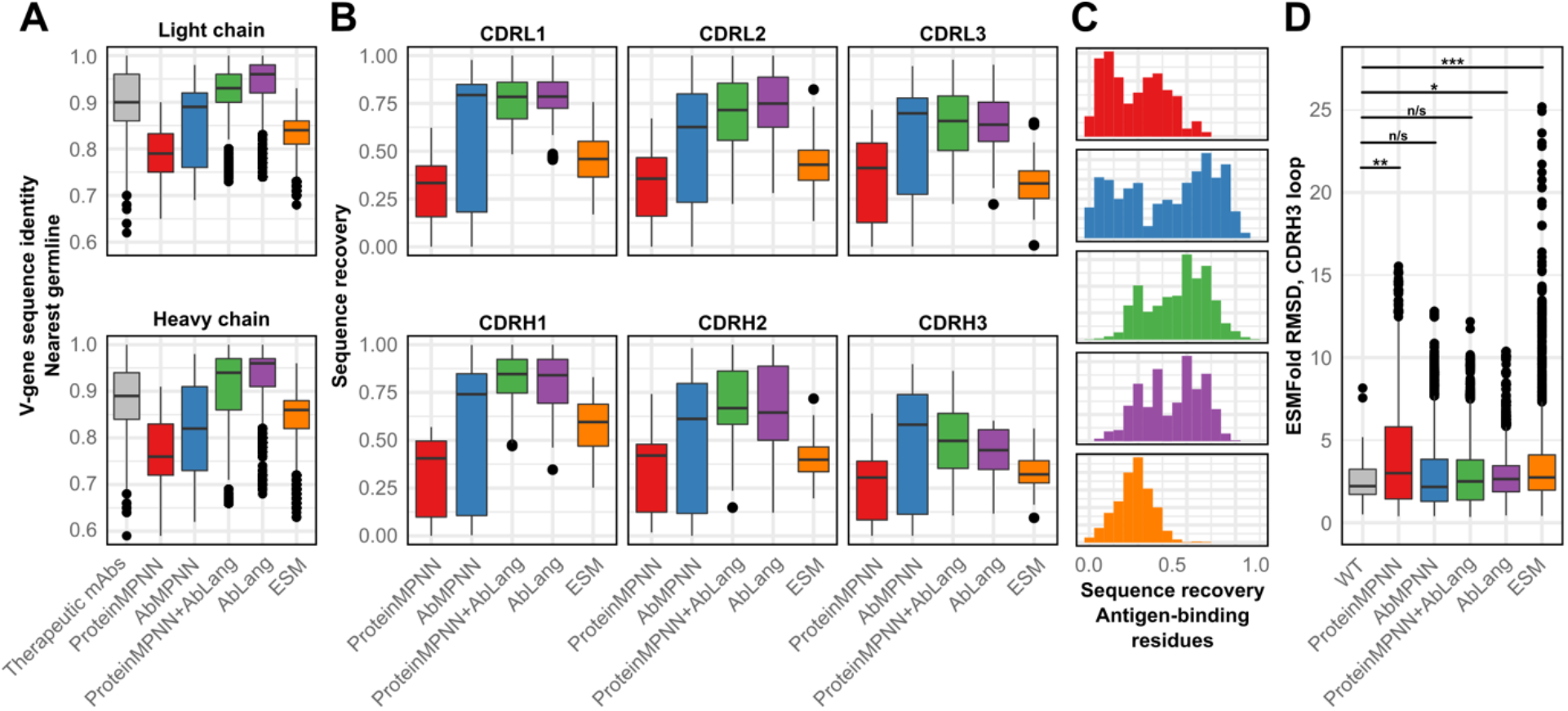
ProteinMPNN does not generate antibody-like sequences unless ensembled with the antibody language model AbLang. **A)** V-gene sequence identity of all designs to the most similar germline. Therapeutic antibody sequences were fetched from TheraSAbDaB on X February 2025. **B)** Recovery of the starting sequence. **C)** Recovery of antigen-binding residues in the CDRs. Y-axis denotes frequency. Top to bottom: ProteinMPNN, AbMPNN, ProteinMPNN+AbLang, AbLang, ESM. **D)** Recovery of the starting conformation of CDRH3. Mann-Whitney U test results: n/s: not significant; *: 0.05≤p<0.5; **: 0.005≤p<0.05; ***: p<0.005.

Second, the designed sequences were generally dissimilar from those of the wildtype antibodies. Poor sequence recovery was observed, consistent with previous reports^16,17^, as measured by identity (i.e., Levenshtein or edit distance, Figure 1B) and chemical similarity (BLOSUM62 dissimilarity matrix, Figure S3). This was especially apparent among antigen-binding residues in the CDRs, with some designs recovering no native residues (Figure 1C), falling markedly short of the reported performance on buried residues of other proteins in the PDB^1^.

Third, the CDR sequences, particularly CDRH3, were inconsistent with structurally similar loops, and were not predicted to refold into the template conformation. Sequence recovery does not reflect the fact that many sequences can refold into the target conformation^6,18^, so the self-consistency of these designs was also evaluated. We compared the designed CDRs to PSSMs derived from PyIgClassify2 clusters, which categorizes them based on their conformations^19^; ProteinMPNN designs were universally less consistent with those PSSMs than designs from other methods (Figure S4). These designs were also compared to structural models predicted by ESMFold^20^, which has been used in previous studies to validate helix- and sheet-rich sequences generated by ProteinMPNN^6,7^. The CDRH3 loops of these models were less similar to the template structure than those of the wildtype sequences (Figures 1D and S5), and the distribution of their RMSDs showed a statistically significant difference from those observed with the wildtype sequences of the benchmark proteins (p=0.003, Mann-Whitney U; antibody-specific structure prediction models^21,22^ were eschewed for this analysis, since they tended to predict an antibody’s CDRs in the same conformation regardless of CDR sequence, even when every CDR residue was mutated to glycine; see Figures S6-S8 for details).

These observations contrasted with designs generated by the antibody language model AbLang, but not the generalist protein language model ESM2^20^. Language models trained on antibody sequences exhibit a strong bias toward germline residues^9^, an observation corroborated here (Figure 1A). Designs generated by AbLang nonetheless exhibited greater sequence recovery, including at antigen-binding sites, as well as lower RMSD to native CDRH3 structures (Figures 1B-1D). The sole advantage observed with ProteinMPNN designs was its capacity to generate diverse sequences for CDRs 1 and 2 (Figure S9).

Given the complementary modalities of these two neural networks, we explored whether including AbLang logits during ProteinMPNN inference could restrain sequence generation toward antibody-like sequences while respecting the structural constraints imposed by the antibody/antigen and heavy/light chain interfaces. The workflow involves jointly designing one random residue at a time by running both neural networks in parallel and adding their logits before softmaxing and sampling (Fig S1). An attractive advantage of this approach is that re-training or fine-tuning is unnecessary. To serve as a reference, a previously published version of ProteinMPNN fine-tuned on antibody models and structures was used (AbMPNN^17^), which also showed marked improvements in many of the metrics discussed above, although its performance was uneven for reasons that were not investigated (Figures 1). However, due to differences in training set composition and the possibility of leakage, AbMPNN was used primarily as a reference point, rather than for head-to-head comparison.

Relative to baseline performance of ProteinMPNN, ProteinMPNN+AbLang showed marked improvement across all metrics, often at similar levels to that of the specialist model AbMPNN. Sequence recovery statistics and refolding RMSD broadly matched the favorable performance of AbLang, particularly in antigen-binding residues (Figure 1C) and the CDRH3 loop (Figure 1D). The V-gene sequence identity of those designs to the nearest germline mirrored values observed in therapeutic antibodies (Figure 1A). Moreover, for CDRs 1 and 2, sequence diversity increased markedly relative to designs made by AbLang only, as measured using the Shannon entropy of the residue compositions at each position (Figure S9). This diversity was not accompanied by an uptick in unreasonable sequences, as judged by the negative log-likelihood values from either PyIgClassify2 clusters or length- and V-gene-family-matched PSSMs (Figures S2 and S3). In short, adding a bias favoring germline residues during structure-based sequence design appears to broadly improve the antibody**-**ness of designed sequences across virtually all metrics assessed here.

To assess the method’s potential to discriminate successful from failed designs, we applied it to an experimental dataset of trastuzumab designs from a previous study^23^. Each method was used to rank the pseudolikelihood values of about 35,000 variants designed by deep mutational scanning^24^. Logits from ProteinMPNN+AbLang outperformed those from either ProteinMPNN or AbLang in isolation when discriminating HER2 binders from non-binders, and nearly matched the performance of the specialist model AbMPNN (Figure 2A). One noteworthy example of how ensembling improves performance is its residue Y105 (Chothia numbering), which has been shown to be indispensable for binding^23,25,26^ and was introduced in 91% of designs from ProteinMPNN+AbLang, 68% from AbLang, and 0% from ProteinMPNN.

**Fig 2.**
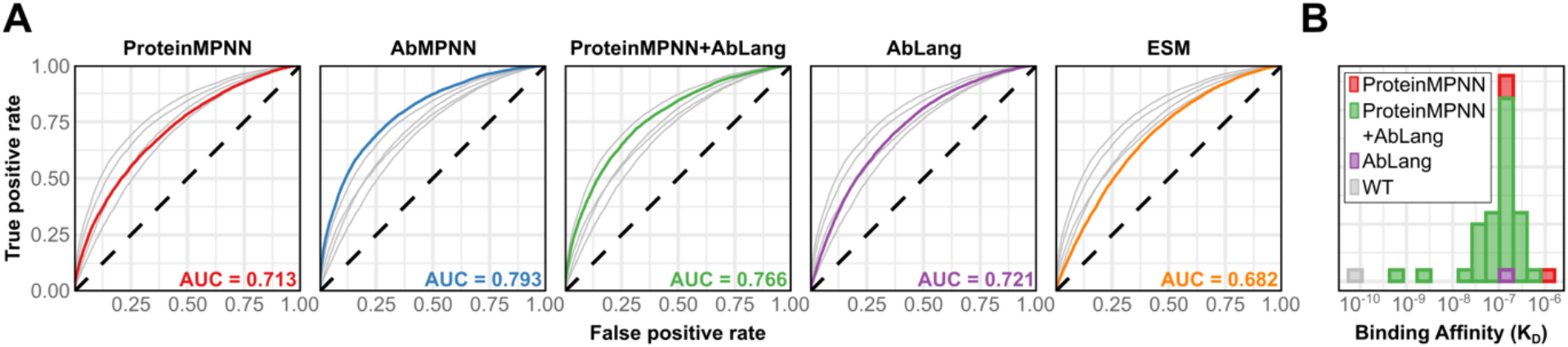
Ensembling ProteinMPNN and AbLang improves identification and design of trastuzumab variants. **A)** ROC curve of ∼35,000 trastuzumab mutants previously measured by deep mutational scanning. Per-residue probabilities were calculated using either the crystal structure with a fully masked CDRH3, or the complete heavy chain Fab (for ESM) or VH (for AbLang) with fully masked CDRH3. **B)** Distribution of experimentally measured HER2 binding affinities of designs generated by ProteinMPNN alone, AbLang alone, or both methods ensembled together. A total of ninety-six designs were generated using each method. Nonbinders were omitted from this chart.

To further evaluate the suitability of ProteinMPNN+AbLang for antibody design, we designed and synthesized 96 trastuzumab CDRH3 variants using sequences designed from ProteinMPNN, AbLang alone, or ProteinMPNN+AbLang. First, one thousand CDRH3 sequences were designed *in silico* using each method. These sequences were clustered into 96 clusters using *k*-medoids^27^, and representatives were selected from each method and expressed in HEK cells (no further filtering was performed). After purification, the antibodies were tested for HER2 binding by surface plasmon resonance (SPR).

The distribution of binding affinities measured by SPR is shown in Figure 2B. Whereas only one AbLang design and three ProteinMPNN designs were found to bind their target, 36 designs from ProteinMPNN+AbLang were found to bind HER2 (p=5e-10 and p=9e-9, respectively, Chi-square test). None of the designs showed greater binding affinity than the wildtype. The overperformance of ProteinMPNN+AbLang in generating binders detected in this experiment reinforces the *in silico* findings detailed above: Constraining ProteinMPNN with a bias toward antibody sequences improves design success rates.

In conclusion, ProteinMPNN’s ability to design well-structured domains does not appear to extend to CDR loops used for antigen binding. Possible reasons for this underperformance include A) lack of relevant training examples in the PDB, relative to structural diversity^28^; B) peculiarities in the sequence composition of antibodies specifically, which evolve somatically from structurally flexible germlines^14,29,30^; and C) misalignments in the training objectives of neural networks for structure-based sequence design, which generate sequences predicted to fold into certain structures without considering whether those sequences do not preferentially adopt alternate conformations^31^. Of these, only the third reason is not ameliorated in some way by AbLang’s inclusion during inference. The language model appears to enforce certain expectations for what naturally occurring antibody sequences ought to look like, for example by limiting ProteinMPNN to the relevant sequence space for CDRs 1 and 2. That said, the strong performance of AbMPNN in some cases, which was trained on several hundred thousand antibody models in addition to the contents of the PDB, shows that inverse folding models should be expected to learn these patterns directly as more structural data become available for training. In the interim, the strategy presented here could serve as a stopgap for designing away from unreasonable antibody sequences, with the advantage that it can be periodically updated as new antibody language models become available^32–34^.

## Methods

### CDR design and analysis

All calculations were carried out using a bespoke version of ProteinMPNN that was refactored for this study. This version also included wrappers for ESM and the heavy and light chain models of AbLang. For each residue to be designed, calculations were carried out on each deep learning model independently and the appropriate logits were added prior to softmax as shown in Figure 1. All designs were carried out using Boltzmann sampling at t=0.1 and executed one residue at a time in a random order (this includes results reported for AbLang only or ESM only, which contrast with their training approach of predicting all masked residues simultaneously). All residues in all CDRs were designed simultaneously.

Antibody structures from RAbD were downloaded from SAbDab in Chothia format^13,35,36^. Three single-chain nanobodies in the original benchmark set were discarded. Non-crystallographic symmetry mates were manually discarded, and the heavy and light chains were renumbered to H and L, respectively, using PyMol^37^. Sequences were then renumbered with AHo/North CDR definitions using ANARCI^38–40^. The antibody structures were then renumbered to AHo using Rosetta Antibody Renumber^13,41^, and minimized using Rosetta FastRelax with Cartesian minimization^42,43^ and the flag `-constrain_relax_to_start_coords`.

One hundred designs were generated using each method on each structure in the benchmark set. BLOSUM62 distance values were calculated using BioPython^44,45^. Negative log-likelihood values against PSSMs from either PyIgClassify2^19^ or a March 2023 download of the Observed Antibody Space database were calculated using an epsilon value of 1e-6. In several cases, the annotated sequence length of a given CDR differed between a PDB structure fetched from SAbDab^35^ and its PyIgClassify2 cluster. Those results for that CDR were not reported in Figure S3. Additionally, cases where fewer than 4000 CDRs from the appropriate V-gene with the appropriate length existed in the Observed Antibody Space were not reported in Figure S2.

### Trastuzumab panel analysis

To compute pseudo-likelihood values from the trastuzumab CDRH3 loop (Figure 2A), all residues in the loop were masked, and predictions for all residues were carried out simultaneously using ProteinMPNN, AbMPNN, ESM, and AbLang-heavy independently as previously described^46^. Then, each sequence was scored by adding the logits of each method:

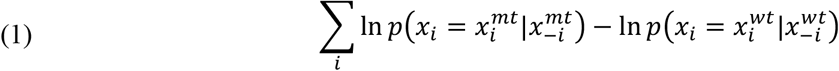

Here,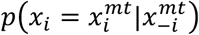 refers to the probability of a mutant with point mutation *mt* at position *i*, given that the unmasked sequence is unmasked everywhere else (*x*_−*i*_). These were normalized to the equivalent value for the wildtype residue. AUROC values were then calculated from these rankings and experimental binding labels using Scikit-learn^47^.

### Trastuzumab variant design, expression, and testing

Trastuzumab variants with redesigned CDRH3 loops were redesigned using the same procedure as above, except only the CDRH3 loop was masked. One thousand designs were generated for each method. The distance between each pair of designs was computed using the BLOSUM62 dissimilarity matrix, and the resulting distance matrix was clustered to 96 clusters using the k-medoids algorithm as implemented in Scikit-learn^27,47^.

Plasmids with designed antibodies were ordered from Twist Biosciences and used to transfect HEK Expi293 cells in Opti-MEM I reduced serum medium with ExpiFectamine Enhancer 1 and ExpiFectamine Enhancer 2. After growing cultures for four days in 1.2 mL, cells were harvested and pelleted by centrifugation at 500g for five minutes, and antibodies were purified using MabSelect SuRe resin eluted with either 0.1M Glycine neutralized with 1 M Tris (pH 7.5), or 0.1M Citrate neutralized with 1 M HEPES (pH 7.6).

SPR experiments were performed using 400 nM HER2 (Acro Biosystems) on a Carterra LSA SPR biosensor equipped with a HC30M chip at 25°C in HBS-TE as previously described^48^. After diluting to 10 µg/mL, antibodies were amine-coupled by EDC/NHS activation to the sensor chip. This was followed by ethanolamine HCl quenching. Analyte was then flowed over the sensor chip in HBS-TE with 0.5 mg/mL BSA with 5 minute and 15 minute association and dissociation, respectively. The surface was regenerated with 2 × 30-second injections of IgG elution buffer (Thermo) following each injection cycle. Carterra’s Kinetics Tool software with 1:1 binding model was used to analyze the data and obtain dissociation constants.

## Acknowledgements

The authors would like to thank Dr. Lewis Chinery for providing the cleaned trastuzumab deep mutational scanning data from Mason & colleagues^23^ that was used in Figure 2A of this manuscript, as well as Drs. Jack Maguire, Jeliazko Jeliazkov, and Tobias Olsen for helpful discussions.

## SUPPLEMENTAL FIGURES

**Figure S1:**
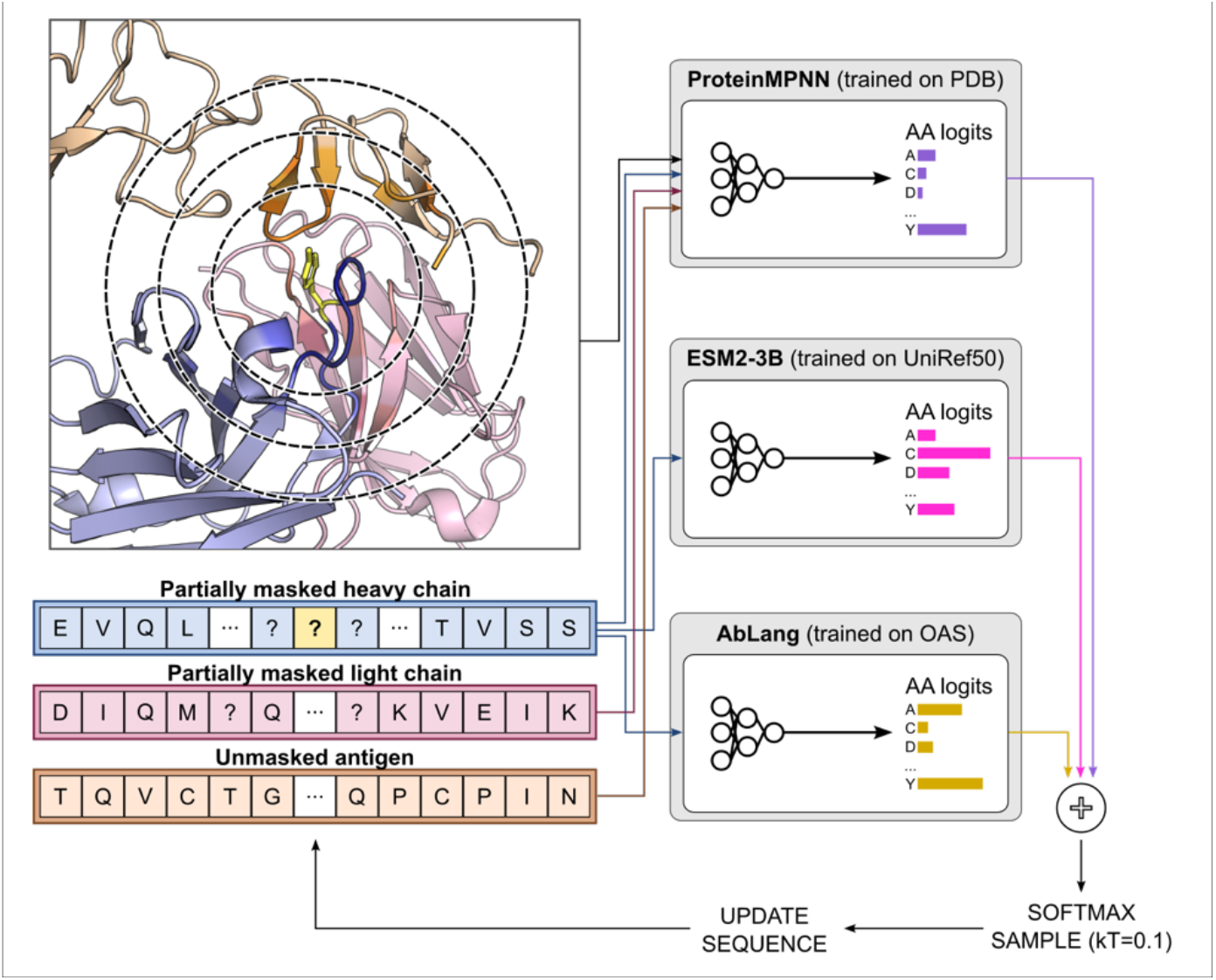
Schematic for ensembling ProteinMPNN, which design residues using local structural characteristics, with protein language models, which do so with global intrachain sequence information.

**Figure S2:**
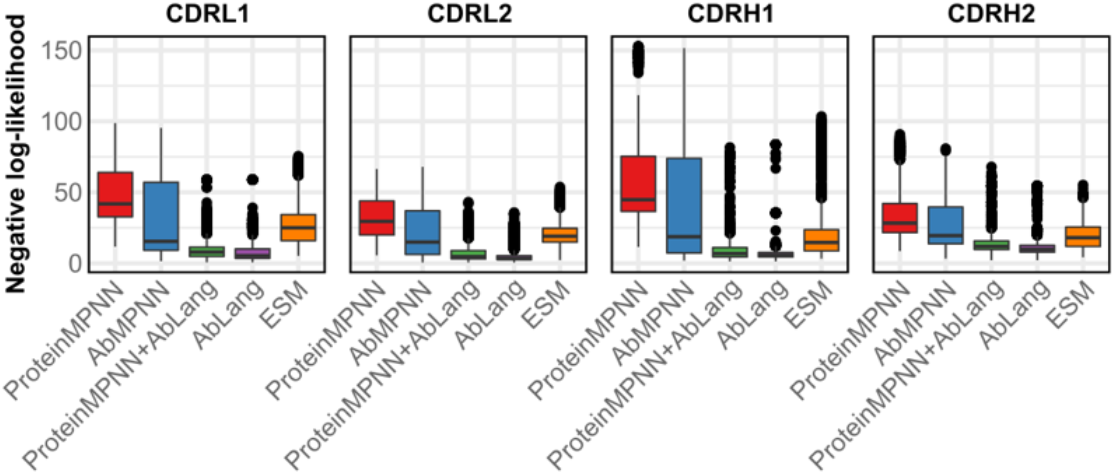
Negative log-likelihood values of CDR designs calculated using V-gene-family- and length-matched position-specific scoring matrices derived from sequences in the OAS.

**Figure S3:**
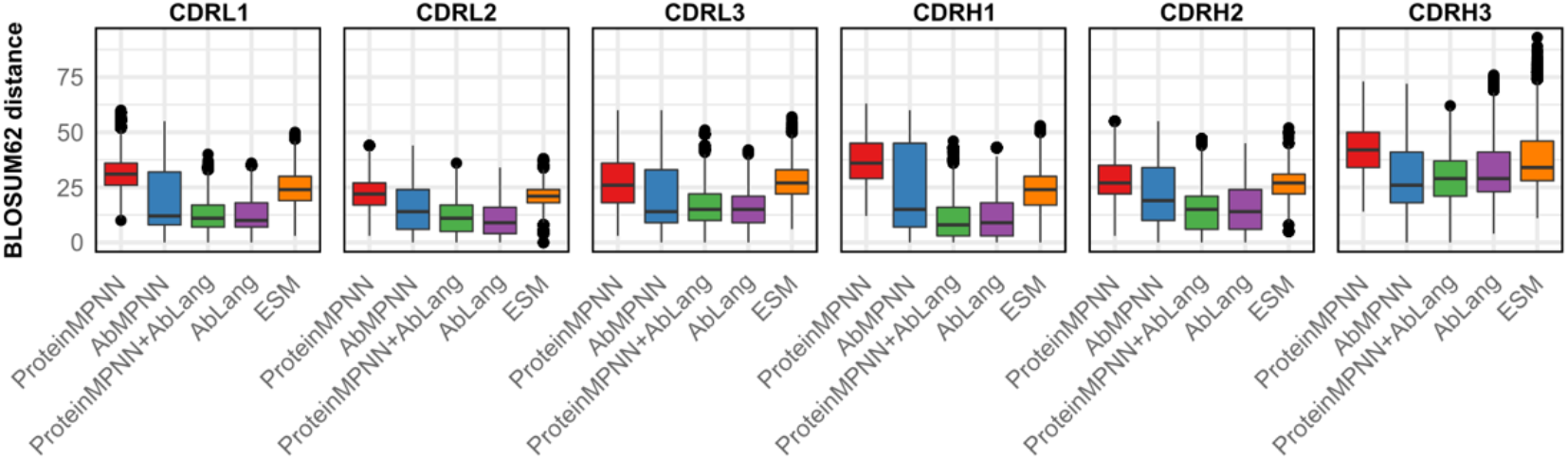
Sequence similarity to WT measured using the BLOSUM62 distance matrix. Lower values indicate greater similarity.

**Fig S4:**
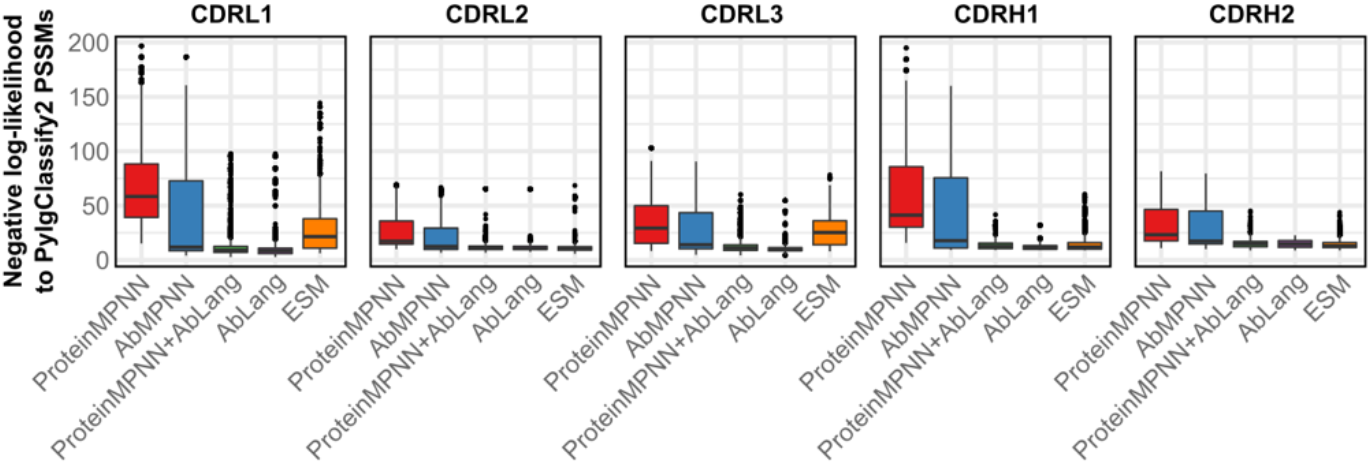
Negative log-likelihood values of sequences measured using PSSMs from PyIgClassify2 conformational clusters.

**Fig S5:**
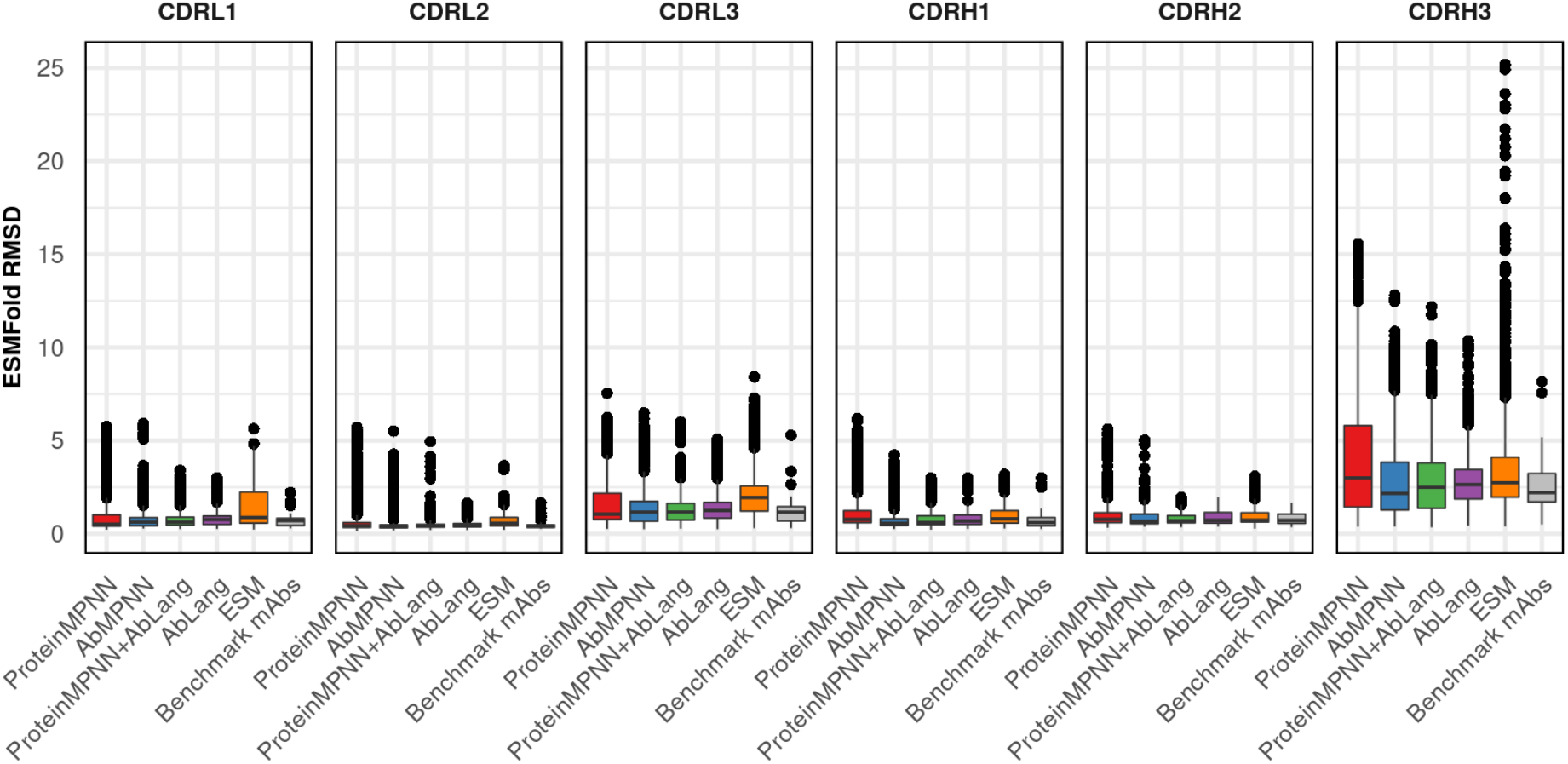
Structure prediction of CDR loops using ESMFold.

**Fig S6.**
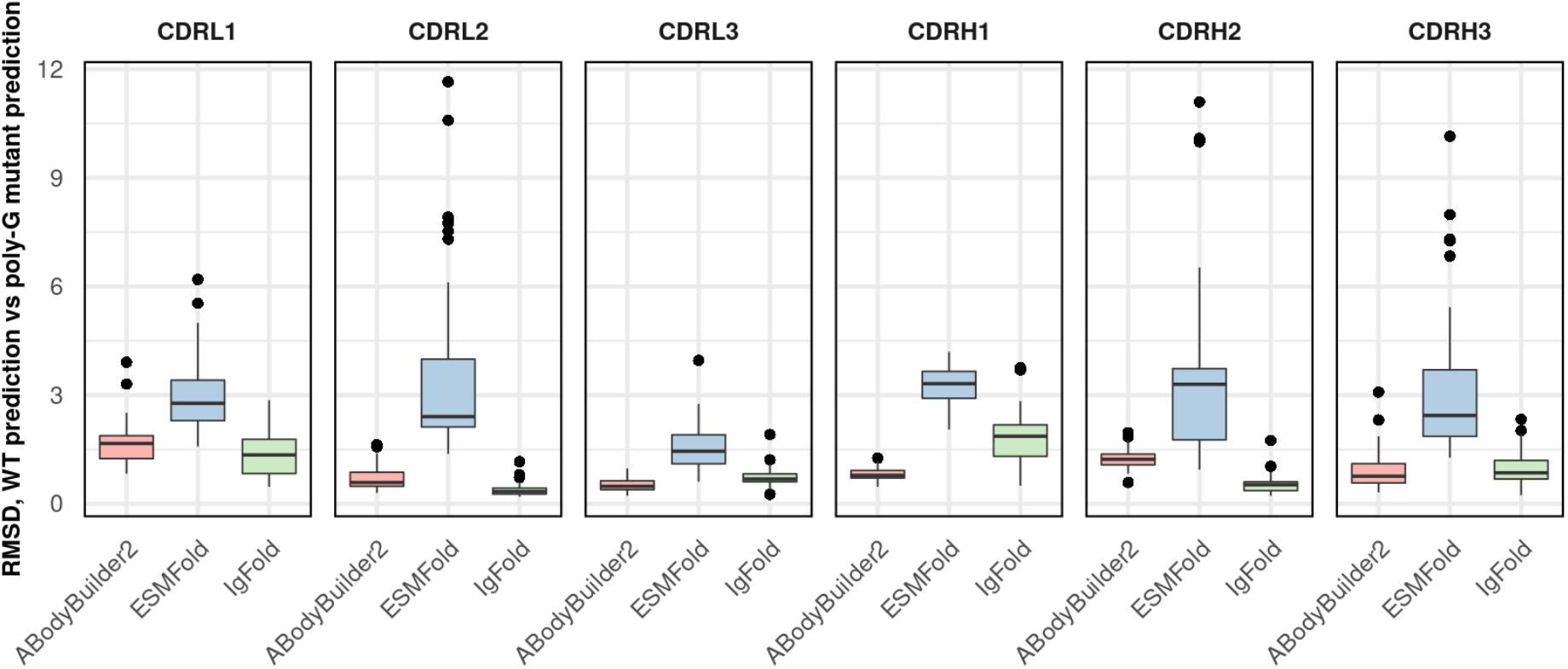
Structure prediction neural networks specific for antibodies return memorized conformations when predicting the conformations of CDRs that have been entirely mutated to polyglycines. In contrast, the general-purpose neural network ESMFold returns perturbed conformations with higher RMSDs. All tested antibodies were part of the training sets of all three neural networks. All CDRs are predicted simultaneously across all tested antibody models, meaning that the mutated sequences have glycines in all six CDRs.

**Figure S7.**
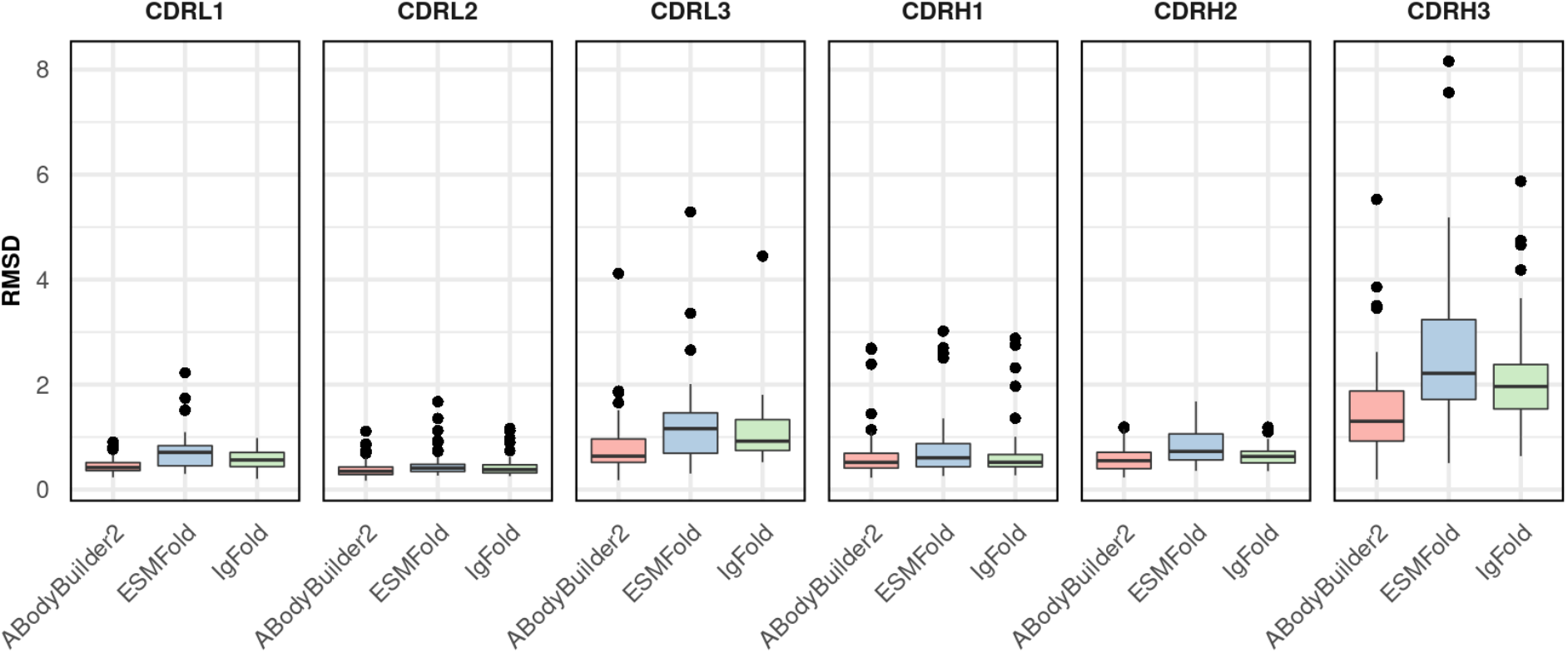
RMSD of CDR loops for antibodies used for design in this study. All tested antibodies were part of the training sets of all three neural networks.

**Figure S8.**
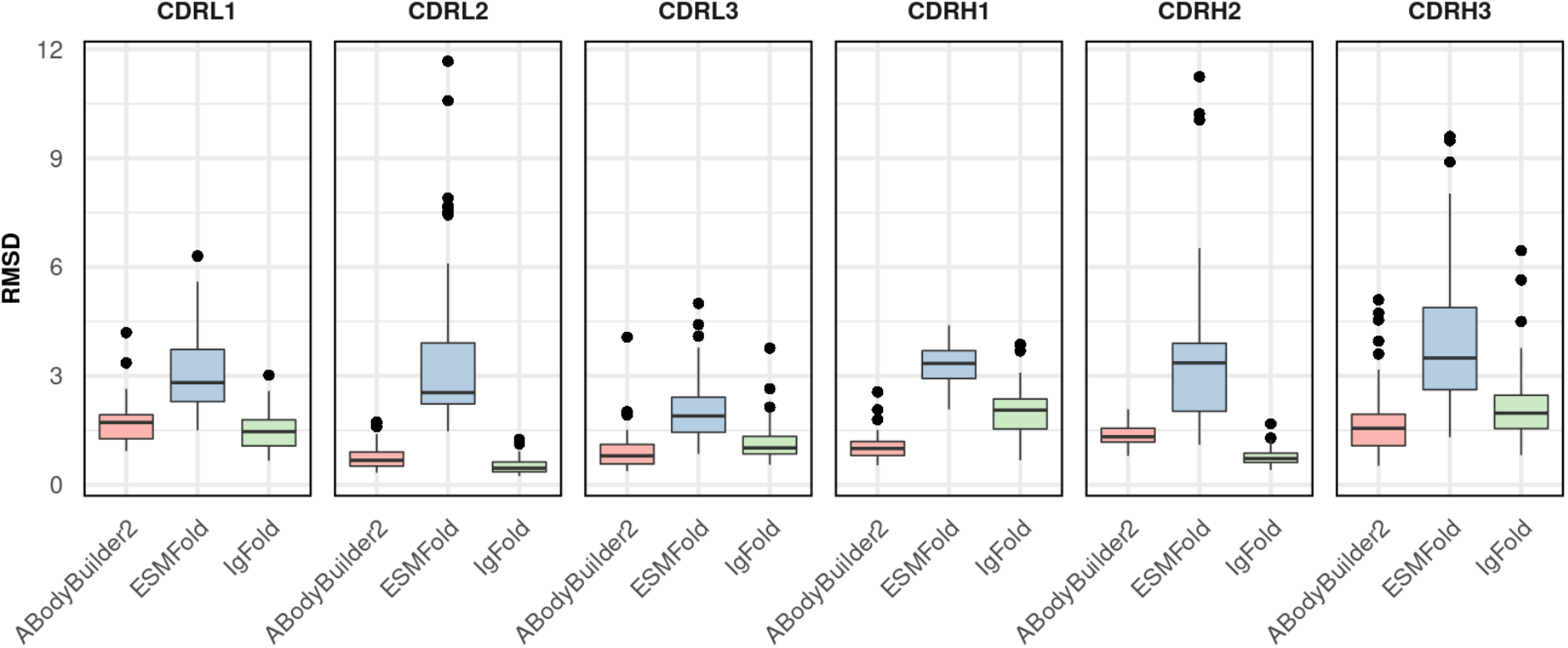
RMSD of CDR loops for antibodies used for design in this study, after mutating every residue in the CDRs to glycine.

**Figure S9.**
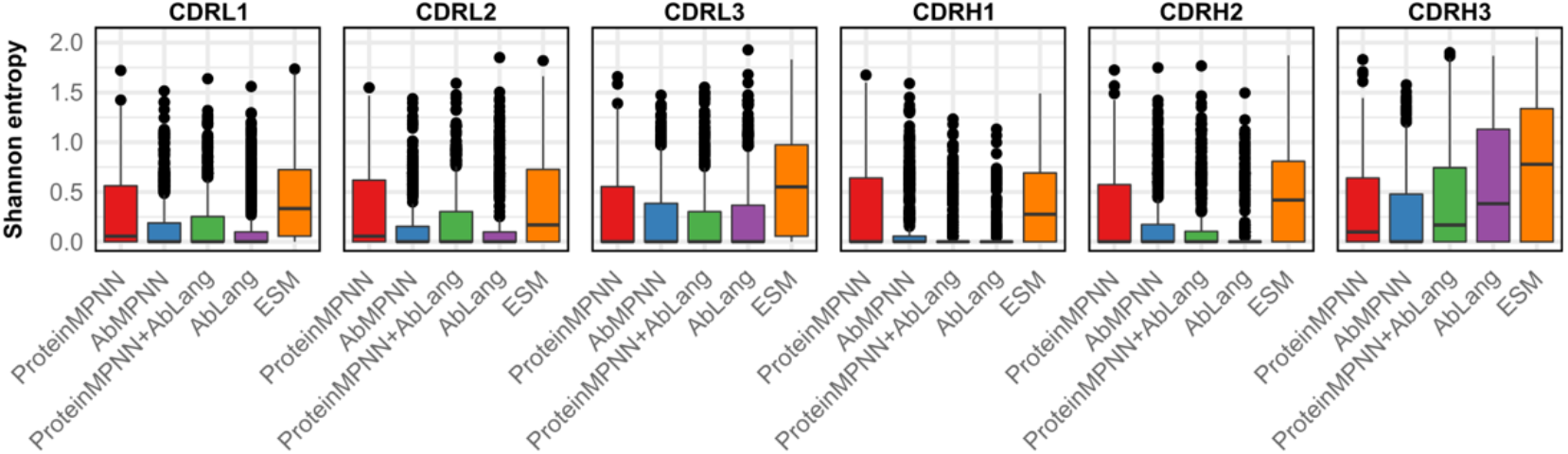
ProteinMPNN designs highly diverse CDRs, as indicated by high Shannon entropy values across different designs.

